# Geographic variation in vulnerability to warming temperatures in an intertidal barnacle species

**DOI:** 10.1101/2024.03.26.586848

**Authors:** Sarah E. Gilman, Gordon T. Ober, Rhiannon L. Rognstad, Madeleine Bunnenberg-Ross, Tingyue Man

## Abstract

Vulnerability to warming temperatures under climate change arises when there is a gap between local climate and local physiology. Intertidal species are unique because they face two distinct thermal environments, and it is unclear which is the bigger driver of thermal physiology and vulnerability. Here we compare the thermal environments and physiology of three populations of the intertidal barnacle *Balanus glandula*, spanning 1460 km of its geographic range. We measured energy consumption in the laboratory across a 5-hour emersion and subsequent 6-hour immersion at 7 different emersion temperatures (10-38°C). We compared these results to one year of emersion and immersion temperature data from each location. Our results suggest that the temperatures experienced during emersion are a bigger driver of each population’s emersion thermal physiology than those experienced during immersion. We also estimated vulnerability to future warming in two ways: as the total annual energy demand and as the number of days above each population’s thermal peak. These produced conflicting results. The central population spent the most days over its thermal peak, but the northernmost population had the greatest total costs over a year. The higher energetic costs in the northernmost population may be explained in part by a strong latitudinal gradient in primary productivity that is selecting for higher energy demand in higher latitude populations. Thus, accurate predictions of *B. glandula*’s response to warming temperatures will require knowledge of both future temperature and food availability.

## 1 Introduction

Identifying the species, and populations within species, most vulnerable to warming temperatures under climate change is a pressing goal of ecology (Helmuth et al. 2002, Bennett et al. 2019). A population’s vulnerability is determined by the magnitude of expected warming relative to its current thermal physiology and to its capacity to alter that physiology through genetic adaptation or phenotypic plasticity. Populations from different locations within a species’ geographic range don’t experience the same temperatures (Helmuth et al. 2002, Deutsch et al. 2008), are unlikely to experience the same amounts of warming (Gilman et al. 2006, Foulk et al. 2024), and likely exhibit different degrees of local adaptation and thermal plasticity (Gunderson et al. 2017, Gaitan-Espitia et al. 2017). This is particularly true for ectotherms, whose body temperatures reflect a complex interaction between the individual organism and its environment (Porter & Gates 1969, Fey et al. 2019, Hayford et al. 2021).

Intertidal habitats present additional challenges to, and opportunities for, understanding inter-population variation in vulnerability to climate change. Intertidal organisms alternate between terrestrial and marine conditions with the daily oscillation of the tides, each with a distinct thermal environment. Because terrestrial climates are more variable and exhibit greater thermal extremes than marine climates, we would expect the two environments to select for very different physiologies. It is generally unknown which of the two environments is the bigger driver of an intertidal organism’s thermal physiology. Many intertidal species are active mainly when immersed and minimize activity and metabolism during low tide (McMahon 1990, Sokolova & Portner 2001, Marshall et al. 2011). Thus natural selection might favor a thermal physiology that is closely matched to the aquatic thermal environment (Sanford 2002, Iles 2014). But intertidal species experience their greatest thermal extremes during emersion and must survive and maintain some metabolic function during those times. Therefore natural selection could also favor physiologies that enhance tolerance and/or performance during emersion. Indeed high shore intertidal species frequently exhibit greater thermal tolerances than their lower shore congeners, despite experiencing the same aquatic thermal environments (Somero 2002, Stillman 2003). Distinguishing the influences of emersion and immersion conditions on population-level thermal physiology requires comparing multiple populations with contrasting emersion and immersion thermal environments.

Approximately 60% of the world’s coastlines experience diurnal or mixed semi-diurnal tides (Byun et al. 2023). In these tidal regimes, aerial exposure is concentrated in a single low or lower-low tide, the timing of which may vary with season, latitude, and coastal geomorphology (Denny 2007). The timing of this low tide can decouple emersion temperatures from latitudinal gradients in terrestrial climate, by filtering the time of day that organisms are exposed to terrestrial conditions (Helmuth et al. 2002, 2006). For example, in the northeastern Pacific, intertidal sites at higher latitudes in Oregon and Washington are much more likely to be exposed to air in middle of the day during the summer than lower latitude locations in California, but the opposite is true in winter (Fig. 1B). This decoupling of emersion and immersion thermal regimes creates a unique opportunity to ask which environment is more important in driving thermal physiology. However, past studies of thermal physiology of NE Pacific intertidal species provide conflicting results. While physiological differences have been identified between populations north and south of a known thermal break at Pt. Conception, CA (Fig. 1A; e.g., Blanchette et al. 2007, Gleason & Burton 2013); intraspecific comparisons at higher latitudes have variously shown associations with low tide temperatures (Sagarin & Somero 2006, Kuo & Sanford 2009), water temperatures (Rao 1953), or patterns uncorrelated with both (Logan et al. 2012).

**Figure 1.**
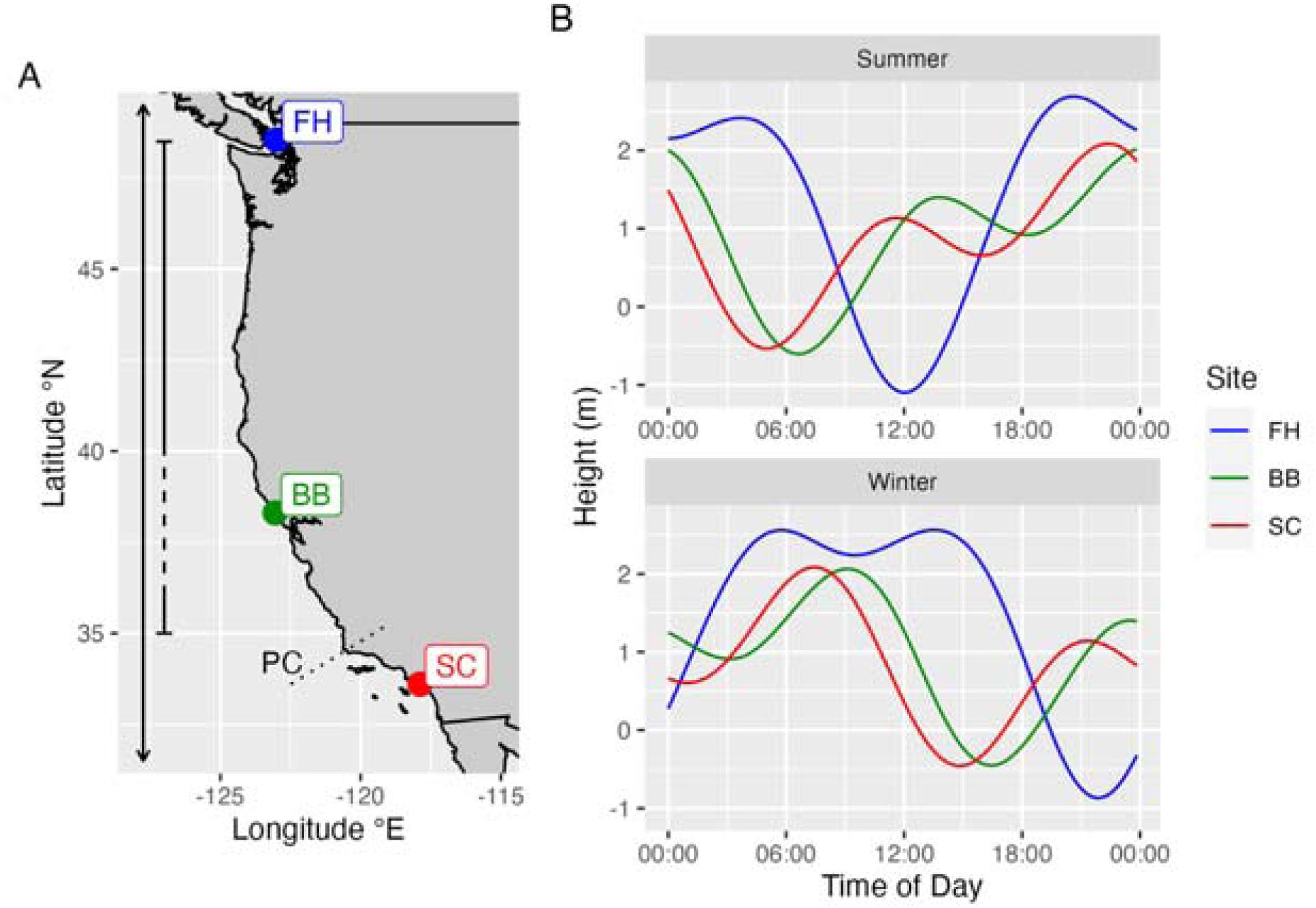
Location of the three populations used in this study (A) and representative summer and winter spring tides for the sites (B). The double-ended arrow in (A) indicates the geographic distribution of *B. glandula*, which extends from Mexico to Alaska (Morris et al. 1980). The flat-headed line indicates the region of genetic sampling in Sotka et al. (2004) and Wares and Skozen (2019), with the genetic cline indicted by the dashed portion. “PC” indicates the thermal break at Point Conception, California. The predicted tides in (B) for June 15, 2022 (“Summer”) and December 23, 2002 (“Winter”) were calculated using the xtide program (http://tbone.biol.sc.edu/tide/index.html)

Here we extend an earlier study of the thermal physiology of the barnacle *Balanus glandula* from single a high latitude northeastern Pacific population (Ober et al. 2019) to two additional sites spanning 1460 km (Fig. 2A) to address two specific research questions: 1) Is terrestrial or marine climate the stronger driver of thermal physiology? and 2) which population is most vulnerable to warming temperatures under climate change? *Balanus glandula* is a common mid to high shore barnacle along the Pacific Coast of North America from Mexico to Alaska (Morris et al. 1980). It has also been introduced to Asia, Africa, and South America (Geller et al. 2008, Simon-Blecher et al. 2008). For each study population, we measured respiration over a 5-hour emersion and a subsequent 6-hour immersion at 7 different emersion temperatures (10, 15, 20, 25, 30, 35, 38°C). We used long-duration trials to fully capture the costs of low tide exposure, as metabolic rates can vary over the duration of a low tide exposure (McGaw et al. 2015, Ober et al. 2019, Griffen et al. 2024).

**Figure 2.**
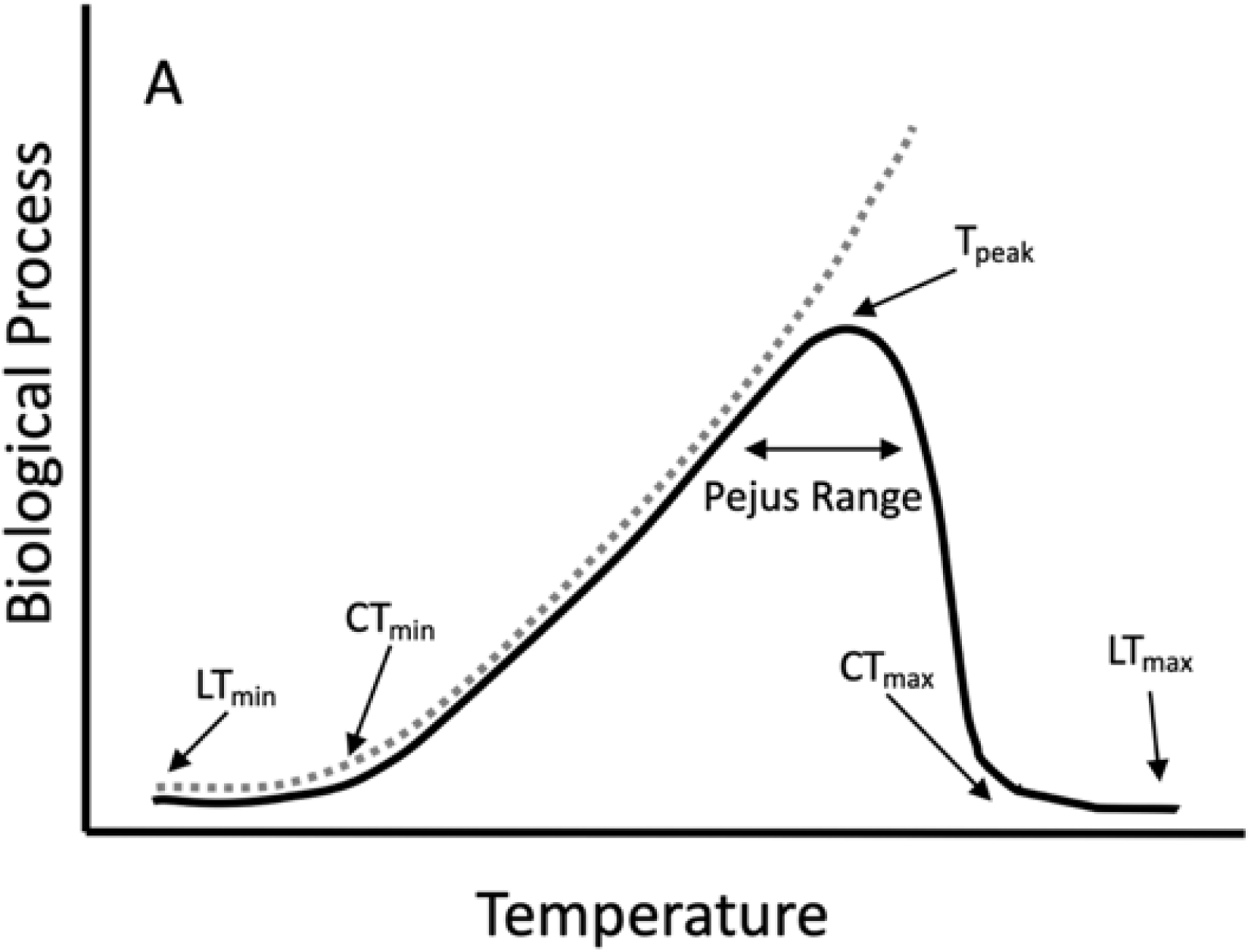
Stereotypical patterns of thermal performance and energetic cost for a poikilotherm. For most traits (solid line), performance increases up to a peak beyond which it decreases. Further warming can lead to reversible (CT_max_) and irreversible (LT_max_) loss of function. Energetic costs of exposure to temperature (dashed line) are usually estimated from a TPC curve for metabolic rate or respiration, with the assumption that the full cost of exposure continues to increase exponentially above T_peak_, even as organism function declines. Figure modified from Hayford et al. (2021).

Based on previous studies (Helmuth et al. 2002, 2006), we expected the aquatic thermal environments of these three sites to correlate with latitude, but the emersion thermal regimes to be uncorrelated. Specifically, we expected the southernmost population, Alamitos Bay (hereafter referred to as SC) to have the warmest immersion and emersion conditions, because it occurs south of Point Conception, CA. We expected the northernmost site, Friday Harbor (hereafter referred to as FH) to have the coldest immersion thermal regime, but an intermediate emersion thermal regime because it had previously been identified as a high latitude hotspot (Helmuth et al. 2002). These three sites also span a known genetic cline between two mitochondrial genotypes in *Balanus glandula* (Wares & Skoczen 2019, Wares et al. 2021). The central site, Bodega Bay (hereafter referred to as BB), occurs in the middle of this cline and the other two sites are beyond its endpoints (Fig. 2A). There is some evidence to suggest that the genetic cline reflects adaptation to different thermal regimes (Wares & Skoczen 2019, Wares et al. 2021).

To assess vulnerability to warming, we compared each population’s thermal physiology to intertidal and water temperature data collected for one year at each study site. Hayford et al. (2021) distinguish two general approaches to assessing a population’s vulnerability to warming under climate change. Discrete or threshold approaches (e.g., Sunday et al. 2019, Bennett et al. 2019, Buckley et al. 2022) quantify the frequency of exposure to a threshold temperature above which organisms experience thermal stress and/or death. Populations that spend more time over such a threshold, either currently or under predicted future temperatures, are considered more vulnerable to warming. Threshold temperatures are usually identified from a thermal performance curve (“TPC”, Fig. 2) of a trait related to fitness. Commonly used thresholds include the critical thermal maximum and lethal limit (Sunday et al. 2010), but the peak of the thermal performance curve (“T_peak_”, also known as the thermal optimum) may be more relevant biologically (Tagliarolo & McQuaid 2015, Buckley et al. 2022) and easier to accurately estimate (Low-Décarie et al. 2017). Continuous approaches (Fly et al. 2012, Montalto et al. 2016) calculate the full energetic cost of exposure to a continuous time series of temperatures for either current or future climates at that location. Populations that show higher total costs under those current or future temperatures are predicted to be more vulnerable to warming. Costs can be estimated from a TPC curve fitted for oxygen consumption, if one assumes that costs continue to increase above T_peak_ even as performance declines (Fig. 2; Sanford 2002, Dell et al. 2011, Lemoine & Burkepile 2012, Ober et al. 2019). Costs can also be incorporated into more detailed energetic models that predict growth rates and/or energy balance (Mislan et al. 2014, Monaco et al. 2014), although this requires knowledge of food availability and intake rates, both of which also may change with temperature (Dell et al. 2011, Englund et al. 2011, Iles 2014). Of the two, the threshold approach is more commonly used because it requires less detailed information, but it has also been criticized for that lack of detail (Buckley et al. 2022). It is not known if the two approaches produce similar predictions.

To address our first question, we estimated the thermal performance curve (Fig. 1) for each population and tested for differences in the peaks of the curves among populations. Because we expected the FH site to have warmer low tide and cooler aquatic conditions than the BB site, we predicted the FH population would have a warmer thermal peak (T_peak_) than BB if emersion temperatures influence physiology but the opposite pattern if water temperature is more important. We also examined temporal patterns of oxygen demand over the duration of individual emersion and immersion trials to test for changes in respiration over time. To address ours second question, we calculated a combined TPC across emersion and immersion phases. We used this curve, in combination with local environmental data, to calculate vulnerability in two ways: the number of days in which temperatures exceeded T_peak_ and the total annual cost of low tide exposure for each population at each site.

## 2 Methods

### 2.1 Study sites and animal collection

The three study sites (Fig. 2) were selected based on their tidal regimes and accessibility for ecological study. All three sites experience mixed semi-diurnal tidal cycles with two unequal low tides on most days. At all three sites animals were collected inside bays or fjords that were protected from most wave action. The northernmost site (“FH”, 48.5457°N, 123.0127W) falls within the Friday Harbor Marine Reserve, managed by the University of Washington, in the Salish Sea. Ober et al. (2019) collected barnacles by deploying 15cm x 15cm PVC settlement plates intertidally off the Friday Harbor Labs dock. The central site (“BB”) was inside the mouth of Bodega Bay CA (“BB”, 38.3037N, 123.0532°W). Here, barnacles were collected both growing on mussel shells within the bay and on PVC settlement plates attached to a rope hung intertidally between pieces of iron rebar installed in the sediment (Quinn et al. 1989). For the southern population (southern California, “SC”), we collected barnacles on oyster shells growing on an artificial jetty on the eastern side of the mouth of Alamitos Bay (33.7484N, 118.1155W).

### 2.2 Temperature Measurements

Intertidal temperature dataloggers were established near each collection site as part of separate study of barnacle growth rates in the field (Gilman et al. *unpubl. data*). For this study we used 1 year of data from loggers located in the approximate vertical center of the barnacle zone. The dataloggers were affixed directly to the rock with Zspar epoxy and recorded temperature every 15-20 minutes. This method directly estimates barnacle temperatures (Wethey 2002, Helmuth et al. 2016), as a large proportion of a barnacle’s surface area is in direct contact with the rock. At FH and BB, we used three replicate Tidbit loggers (v1 or v2 Onset Computer Corp., Bourne, MA, USA), spaced several meters apart. The FH datalogger site was on a west-facing rock wall, approximately 100 m from the collection site. The BB datalogger site was on the jetty inside the mouth of Bodega Bay and within the area used to collect barnacles. For SC, two replicate ibutton dataloggers (DS-1921G or DS-1922L, Analog Devices Inc., Wilmington MA, USA) were placed on a southeast-facing artificial sea wall in Newport Bay (33.6022°N,117.8824°W), approximately 27 km from the collecting site. Loggers were downloaded and replaced every 6 months at BB and FH and every 2-4 weeks at SC.

Water temperature data was downloaded from established monitoring stations near each site. The FH site contains a NOAA monitoring station (9449880, https://tidesandcurrents.noaa.gov/stationhome.html?id=9449880). Water temperature data for BB came from the Bodega Ocean Observing Node (https://boon.ucdavis.edu/data-access/products) approximately 2km from the collecting site. Water temperature data for SC came from the SCCOS Automated Shore station at Newport Pier (https://erddap.sccoos.org/erddap/index.html), approximately 1.7 km from the sea wall with the intertidal dataloggers.

We calculated daily physiological maxima (DPM) for the intertidal and water temperature datasets as the 97.9 percentile of the daily data, after first selecting the maximum temperature among any replicate dataloggers within each site and timepoint. This percentile reflects a temperature exposure of at least 30 minutes and is more physiologically relevant than the absolute daily maximum, which may only be experienced for a minute (Helmuth & Hofmann 2001, Fitzhenry et al. 2004, Gilman et al. 2006, Helmuth et al. 2006).

### 2.3 Laboratory acclimation conditions

As described in Ober et al. (2019), collected barnacles were isolated, tagged, and their maximum opercular diameter measured. Barnacles were maintained in the laboratory in a tidally cycling tank with a daily light cycle (Table 1). They were fed brine shrimp (*Artemia spp.*, Pentair Aquatic Eco-Systems, Apopka, FL 32703) two to three times per week (Gilman et al. 2013). Laboratory tidal regimes (Table 1) were based on local tidal cycles for the warmest part of the year at each site (approximately July-October for SC and BB and April – August for FH), at a height equivalent to the middle of the barnacle zone. The acclimation water temperatures were calculated by averaging at least one year of water temperature data from the local water monitoring station. We acclimated each population to its local average water temp, rather than a common water temperature amongst the three populations, in order to capture ecologically relevant responses to the emersion treatments. Additionally, there is little overlap in water temperatures between SC and the other two sites (Fig. 3), meaning a common water temperatures would be stressful to at least one population. Animals were acclimated to the conditions for at least a week before each respiration trial and re-acclimated for at least one week between trials. They were starved for 48 hours prior to each trial.

**Figure 3.**
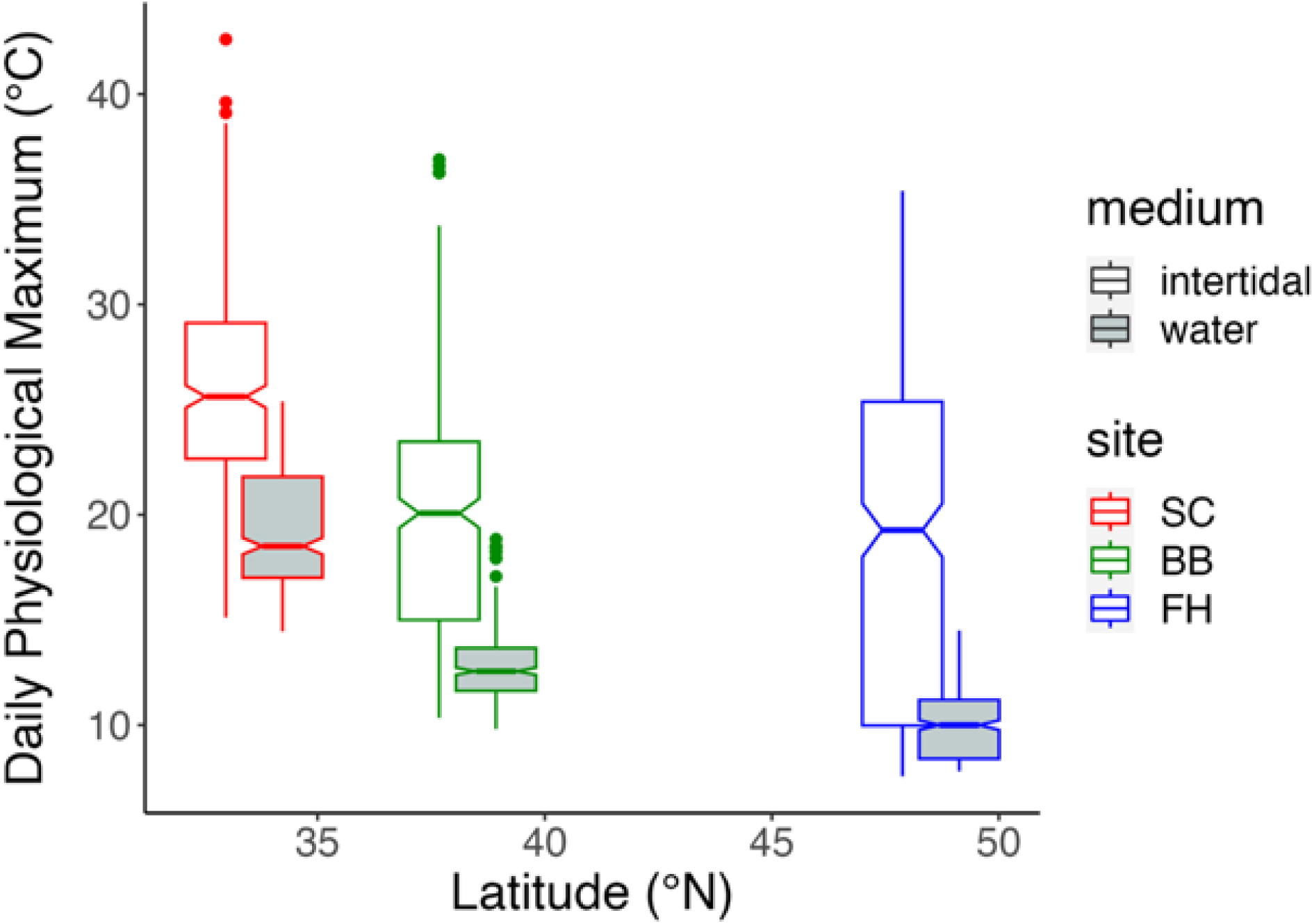
Box plot of daily 97.9%ile of intertidal data logger and water temperatures at each site over one year. The top and bottom of the box indicate the 25th and 75th percentiles. The notched line is the median. The notch breadth indicates 95% CIs around the median.

**Table 1.** Laboratory tidal cycles and abiotic conditions used for each population.

|  | **Friday Harbor, WA** | **Bodega Bay, CA** | **Southern California** |
| --- | --- | --- | --- |
| **Low Tide 1** |  | 7:30am – 12:30 pm |  |
| **High Tide 1** |  | 12:30pm – 7:30pm |  |
| **Low Tide 2** | 7:30pm – 10:30pm | 7:30pm-3:30am | 7:30pm – 1:30am |
| **High Tide 2** | 10:30pm – 7:30am | 3:30am-7:30am | 1:30am – 7:30 am |
| **Lights on** | 03:30am – 7:30pm | 04:45am – 07:00pm |  |
| **Water Temperature** | 10°C | 12°C | 18°C |
| **Air Temperature** |  | 18-20 °C |  |

### 2.4 Respiration experiment

Full methods for the respiration protocol are reported in Ober et al. (2019). Briefly, about 30 minutes before the end of the “High Tide 2” phase of the laboratory tidal cycle (Table 1), barnacles were placed into individual 20ml glass chambers filled with filtered seawater. These were submerged in a temperature-controlled water bath at the acclimation water temperature. After 30 minutes, the water was siphoned out of the chambers and the water bath was heated or cooled until the interior of the chambers reached the experimental temperature. They remained at the experimental temperature for the remainder of the 5-hr low tide period. Oxygen concentrations in the chambers were recorded every 10-seconds with a fluorometric oxygen system with non-invasive sensor spots (Fibox 4 with Pst3-NAU spots, PreSens Precision Sensing, Regensburg, Germany). An empty chamber was included as a control. Afterwards, the barnacles were transferred to individual aquatic respiration chambers and submerged in a second water bath at the population’s acclimation water temperature. Aquatic respiration was monitored for 6 hours.

While Ober et al. (2019) used a fixed ramping rate of 10°C hr-1 for the FH population, this study used a fixed ramping period of 2.5 hrs. To facilitate comparisons among sites, we only used emersion data from the last 2.5 hours of each emersion trial, when all trials were at the experimental temperature. Additionally, in the present study we used 8 blocks of 4 barnacles from SC and BB, with each block used at two different temperatures, for a total of 8 barnacles per temperatures. The FH experiment used 9 barnacles per temperature, in blocks of 3.

### 2.5 Calculation of respiration rates

Slopes of change in oxygen concentration were calculated every five minutes for the last 2.5 hours of the emersion phase and the full six hours of the immersion phase. Slopes were only calculated for intervals with at least 10 data points. We divided the emersion observations by the control value before calculating the slope, in order to correct for the effect of pressure changes on the oxygen readings within the vials during the ramping phase. For aquatic data, we subtracted the slope of the control chamber from each experimental chamber to control for backgrounds rates of oxygen consumption within the seawater. If a control slope was missing for a five-minute aquatic interval, we substituted the average control slope for that trial. We used the five-minute slopes to calculate mean respiration rates for every 30 minutes of the last 2.5 hours of the emersion phase and for every hour of the immersion phase. For this we used 10% trimmed means to systematically exclude outliers. All calculations were completed using the Presens Oxygen Calculator (v 2.2.6) and R (v 4.2.2).

### 2.6 Statistical analyses

All statistical analyses were done in R 4.2.2, using the packages lme4 v. 3.1.160 (Bates et al. 2015), nlme v. 1.1.31 (Pinheiro et al. 2022), car 3.1-1(Fox & Weisberg 2019), and emmeans 1.8.3 (Lenth 2022). In all models that included size as covariate, we used (Operculum Length)^1.8533^, based on a previously reported allometric relationship (Gilman et al. 2013). We adjusted for multiple comparisons among sites using the Tukey method. We used the deltaMethod function of the car package to compare nonlinear parameter combinations.

To compare the intertidal and water DPMs, we used analysis of variance with autocorrelated errors in the gls function of nlme. We also compared the 25%, 50%, and 75%ile of the intertidal temperatures using the rq function of the quantreg package v. 5.94 (Koenker 2022).

All respirometry data were analyzed in mixed models following Zuur et al (2009). First a full model was built, with all possible random and fixed terms. The random terms, which included trial date, individual barnacle, and block, were selected using AIC and this model was tested for departures from normality and heteroskedasticity. The fixed effects were then selected in backwards stepwise approach, starting with the highest order interaction terms, with type II likelihood ratio χ^2^ tests. Normality and heteroskedasticity were re-tested in this final model.

To compare thermal physiology thresholds among populations, we fit quadratic thermal performance curves in each medium and compared the location of T_peak_. We first averaged across the hourly or half-hourly time points to calculate an average respiration rate per barnacle. We then estimated the thermal performance curves for each population in each medium as a parabola of the form: µmol O_2_ = ß_1,P_ + ß_2,P_T + ß_3,P_T^2^ + ß_4_Op^1.8533^, where P=source population, T=temperature, and Op=operculum length. From this we calculated T_peak_ for each population as 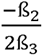. We selected this simple quadratic curve as curves with fewer parameters are recommended when there are a small number of study temperatures (Low-Décarie et al. 2017). We included only barnacle identity as a random factor in both the emersion and immersion models, as it had better AIC values than any other combination of random terms. Due to significant error heteroskedasticity, both models included extra error terms to allow for heterogeneous variances among temperature treatments (immersion) or temperatures and source populations (emersion).

The long duration of the trials also allowed us to look for temporal pattern in respiration rates. For example, a decreasing emersion respiration rate over time could suggest physiological impairment, while the same pattern during immersion would be consistent with an oxygen debt. We tested for temporal patterns by fitting a separate mixed-model ANCOVA for each medium. These models treated temperature (T) as a categorical variable and also included source population (P) and time point (H) as categorical factors and Operculum Length (Op) as a covariate. They were of the form: µmol O_2_ = ß_1_ + ß_2_P + ß_3_H + ß_4_T + ß_5_PH + ß_6_PT + ß_7_HT + ß_8_PHT + ß_9_Op^1.8533^. We included only barnacle identity as a random factor in both the emersion and immersion models, as it had better AIC values than other combinations of random terms. Due to significant error heteroskedasticity, both models included an extra error term to allow heterogeneous variances among trial dates. One data point for an FH barnacle at 38°C at the 30-minute point was excluded from the emersion data set, as it was 5-fold larger than any other data point. Polynomial contrasts were used to identify respiration patterns over time within each combination of source population and experimental temperature.

### 2.7 Vulnerability to warming

To estimate each population’s vulnerability to warming temperatures, we first calculated the cost of a 30-minute emersion at each temperature for each phase of the trial using the estimated marginal means (EMM) from the quadratic models described above. For the cost during emersion, we multiplied the 5-minute EMM values from the emersion model by 6 to scale up to 30 minutes. For immersion, we subtracted off a baseline rate of oxygen consumption from the 5-minute immersion EMMs, using the EMM for hour 6 of from the temporal analysis as the baseline, and then multiplied by 72 to get the full oxygen debt for the 6-hr immersion trial. We divided this value by 5 to adjust it from the 2.5-hour emersion exposure of the trial to a 30-emersion exposure. Finally, we summed the emersion and immersion values to get the total oxygen demand of a 30-minute emersion at each temperature.

We used these values to calculate vulnerability in two ways: as the number of days spent above T_peak_ and as the total costs of emersion over one year. In the first case, we fit a new TPC parabola for each population from the total cost values and calculated T_peak_ as described above. In the second case, because costs increase with temperature even though TPC curves decline at high temperatures declines (Fig. 1; Sanford 2002, Dell et al. 2011, Lemoine & Burkepile 2012, Ober et al. 2019), we calculated an exponentially increasing total cost function based on the metabolic theory of ecology equation (Brown et al. 2004, Dell et al. 2011, Ober et al. 2019), excluding temperatures above the peak of the total cost curve (or below the minimum for SC).

## 3 Results

### 3.1 Field temperatures

There was clear pattern of declining water temperatures with increasing latitude, but no such pattern for intertidal temperatures (Fig. 3). Both intertidal (F_2,1093_ = 21.8, p<0.0001) and water (F_2,1093_ = 38.02, p<0.0001) DPMs showed significant differences among sites, driven mainly by the southernmost SC site which was significantly warmer than BB and FH in both mediums (p < 0.001 in all comparisons). The mean intertidal DPM at the SC site (26.0, CI: 23.7-28.4°C) exceed both the BB (19.5, CI:17.5-22.2°C) and FH (18.4, CI:16.0-20.7°C) sites by more than 6°C. The mean water DPM for SC (19.8, CI: 17.9-21.7°C) was similarly more than 6°C warmer than the mean water DPMs at BB (12.9, CI:10.0-14.8°C) and FH (9.8, CI:7.8-11.7°C). The northernmost FH site had a 3°C cooler mean water DPM and a 1.5°C warmer mean intertidal DPM than the BB site, but these differences were not significant (p >0.05). Similarly, the quantile regression revealed no significant difference in the median intertidal DPM between BB and FH (p=0.634). However, all three sites differed significantly in their the 25^th^ and 75^th^ percentiles of the intertidal DPMs (p<0.004 for all comparisons). The southernmost (SC) site had both the warmest 75%ile and 25%ile for intertidal temperatures, while the northernmost (FH) site had the coldest 25%ile and the central BB site had the coldest 75%ile (Fig. 3). Thus, the FH site had the greatest range of intertidal DPMs (Fig. 3).

### 3.2 Thermal performance curves

When respiration rates were pooled over timepoints, both the immersion and emersion analyses revealed a statistically significant quadratic term, indicating that parabolas fit the respiration data better than linear regression (Table 2A). There was a significant main effect of population, with FH showing significantly higher respiration than SC or BB at most temperatures (p<0.05). There were also significant interactions between population and both the linear and quadratic temperature terms, indicating differences in the shape or position of the parabolas among the three populations (Table 2A, Fig. 4). The fitted parabolas (Table 2B) for emersion for the two most northern populations were concave, with T_peak_ estimated at 27.5°C (23.8-31.2 95%CI) for FH and 24.9°C (19.2-30.5 95%CI) for BB (Fig. 4a and 4c). In contrast, the parabola for SC (Table 2B) was convex (Fig. 4e), with a minimum respiration rate at 20.0°C (14.4-25.6 95%CI). Multiple comparisons tests revealed a significant difference in the location between the emersion peaks for FH and SC (p=0.028), but no other differences were significant (p>0.05).

**Figure 4.**
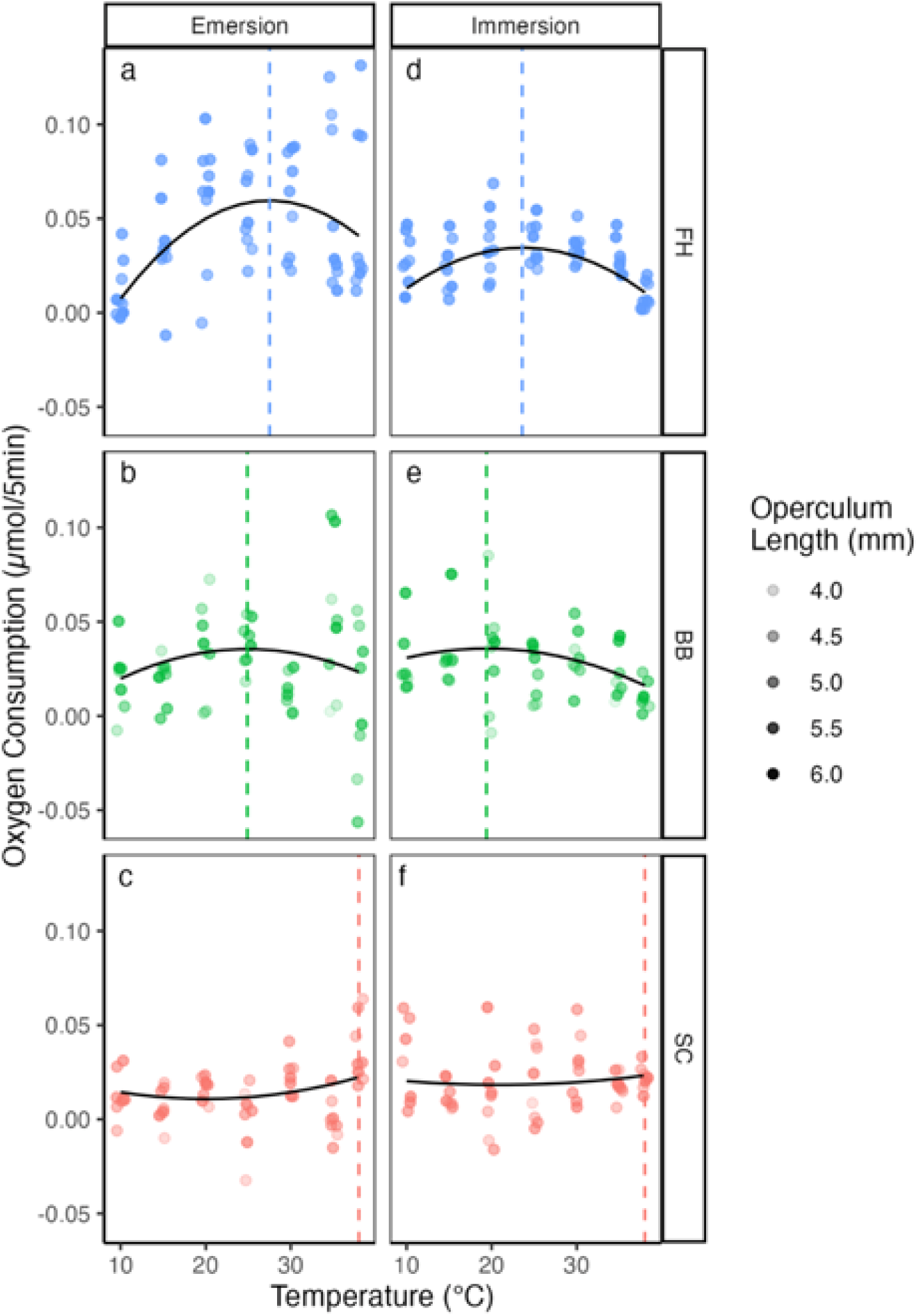
Oxygen consumption by temperature for each population during emersion (A-C) and immersion (D-F) phases. Parabolas represent the predicted respiration rates from the mixed model, calculated at the mean overall barnacle size (5.05 mm operculum length). Dashed lines are the estimated T_peak_ values. The individual points are mean observed respiration rates for individual barnacles, with the darkness of the fill indicating size.

**Table 2.** Model results for average oxygen consumption, treating temperature as a continuous variable. . Analysis of deviance statistics for each medium (A) and parameter estimates (+se) for the line of best fit from the model for each population (B).

| A) | Emersion |  |  | Immersion |  |  |
| --- | --- | --- | --- | --- | --- | --- |
| Term | $\chi^2$ | df | p | $\chi^2$ | df | p |
| Population | 32.5011 | 2 | <0.0001 | 3.2237 | 2 | 0.1995 |
| Temperature | 2.5823 | 1 | 0.1081 | 13.0248 | 1 | 0.0003 |
| Temperature<sup>2</sup> | 0.8269 | 1 | 0.3632 | 16.6997 | 1 | < 0.0001 |
| Size | 5.1147 | 1 | 0.0237 | 12.8529 | 1 | 0.0003 |
| Pop x Temp | 26.224 | 2 | <0.0001 | 13.5512 | 2 | 0.0011 |
| Pop x Temp<sup>2</sup> | 32.5011 | 2 | <0.0001 | 16.1523 | 2 | 0.0003 |

| B) | Bodega Bay | Friday Harbor | Southern California |
| --- | --- | --- | --- |
| Emersion |  |  |  |
| Intercept | -0.03280 (0.02094) | -0.09348 (0.0235) | 0.00059 (0.01494) |
| Temperature | 0.00122 (0.00056) | 0.00353 (0.00185) | 0.00935 (0.00201) |
| Temperature^2 | -0.00142 (0.00101) | -0.00007 (0.00004) | -0.00017 (0.00005) |
| Operculum Length | 0.00004 (0.00002) | 0.00004 (0.00002) | 0.00004 (0.00002) |
| Immersion |  |  |  |
| Intercept | -0.01169 (0.01698) | -0.05631 (0.01585) | -0.00053 (0.01652) |
| Temperature | 0.0022 (0.00138) | 0.00546 (0.00117) | -0.00071 (0.00132) |
| Temperature^2 | -0.00006 (0.00003) | -0.00012 (0.00002) | 0.00002 (0.00003) |
| Operculum Length | 0.00131 (0.00038) | 0.00131 (0.00038) | 0.00131 (0.00038) |

A similar pattern was seen in the immersion curves (Table 2B). The two most northern populations were concave, with T_peak_ estimated at 23.6°C (22.3-25.0 95%CI) for FH and 19.4 °C (13.2-25.5 95%CI) for BB (Fig. 4b and 4d). But the southern site (SC) again showed a convex pattern (Fig. 4f), with a minimum respiration rate at 20.9°C (4.9 – 36.9 %CI). These differences were not statistically significant (p>0.05).

### 3.3 Temporal patterns of respiration by medium

#### 3.3.1 Emersion

The emersion respiration rates ranged from 0.0018 to 0.057 µmol/5min, depending on the source population, temperature, and time point (Fig. 5). All terms in the mixed model ANCOVA for emersion were statistically significant, including the three-way interaction between source population, time, and treatment temperature (Table 3). Most combinations of source population and temperature showed little variation across timepoints, with two exceptions. First, within the southernmost (SC) population, all but the two lowest and one highest temperature showed a significantly increasing linear trend in respiration rates over the 2.5 hr. period. Second, above 20 °C, both BB and FH showed linearly decreasing trends over time.

**Figure 5.**
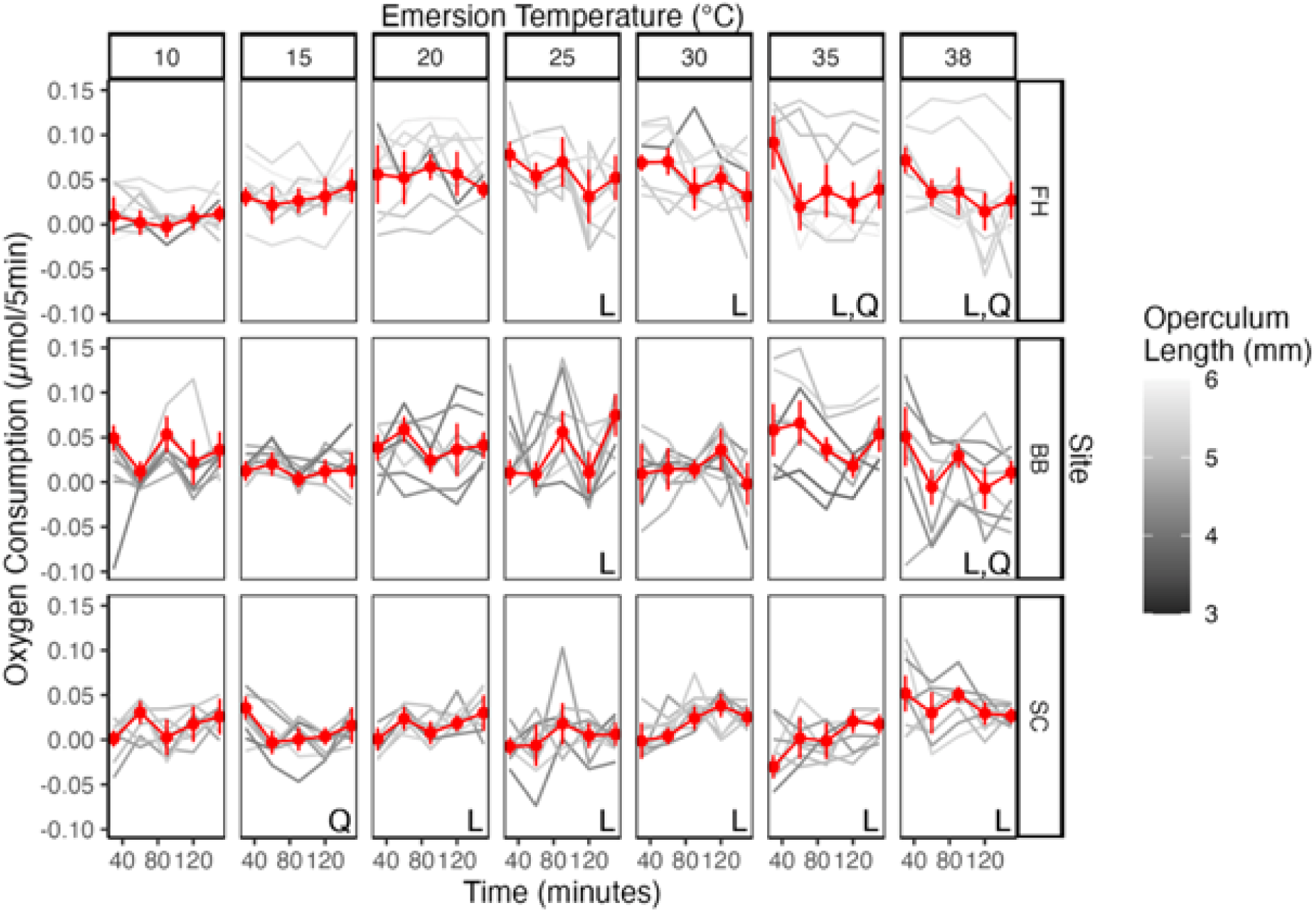
Average emersion oxygen consumption by temperature and timepoint for each population. Gray lines represent data for individual barnacles, darkness reflects size. Estimated marginal means (±95CI), for a barnacle of 5.05 mm operculum length, are shown in red. Letters indicate significant polynomial contrasts (p<0.05, L=Linear, Q=Quadratic)

**Table 3.** Analysis of deviance for respiration by population and timepoint during emersion and immersion.

| **Term** | **Emersion** |  |  | **Immersion** |  |  |
| --- | --- | --- | --- | --- | --- | --- |
| | $\chi^2$ | df | p | $\chi^2$ | df | p |
| **Population** | 36.6454 | 2 | <0.0001 | 4.8558 | 2 | 0.0882 |
| **Time** | 7.7978 | 4 | 0.0993 | 76.7112 | 5 | <0.0001 |
| **Temperature** | 204.9101 | 6 | <0.0001 | 288.4884 | 6 | <0.0001 |
| **Size** | 4.5776 | 1 | 0.0324 | 8.8019 | 1 | 0.0030 |
| **Pop x Time** | 71.1303 | 8 | <0.0001 | 10.7962 | 10 | 0.3736 |
| **Pop x Temp** | 237.3474 | 12 | <0.0001 | 349.9856 | 12 | <0.0001 |
| **Temp x Time** | 235.9039 | 24 | <0.0001 | 97.2488 | 30 | 0.0000 |
| **Pop x Time x Temp** | 294.976 | 48 | <0.0001 | 128.1517 | 60 | 0.0000 |

#### 3.3.2 Immersion

Average immersion respiration rates ranged from 0.0071 to 0.0431 µmol/5min, depending on the source population, temperature, and time point (Fig. 6). The mixed model analysis revealed a significant overall effect of emersion temperature on immersion respiration, as well as significant differences among populations in the form of interactions between population and temperature and between population, temperature, and time (Table 3). There were three distinct temporal patterns among the temperatures (Fig. 6). At the coolest temperatures we most frequently detected a significant quadratic trend in each population, where immersion respiration rates tended to be lowest in the first hour post emersion, peak in the second or third hour, and then decline. This pattern occurred at temperatures between 10-20°C at the central BB site, between 10-25°C at the northern FH site and at temperatures up to 30°C at the southern SC site, although the quadratic trend was not always statistically significant (Fig. 6). Above these temperatures, the pattern shifted to a linear negative trend, indicating higher respirations rates in the first 1-2 hours post emersion. This occurred at 25 and 35°C for BB, 30 and 35°C for FH and 35°C for SC. Finally, at 35 and 38°C for BB and at 38°C for the other two sites, respiration was initially depressed and then increased over time.

**Figure 6.**
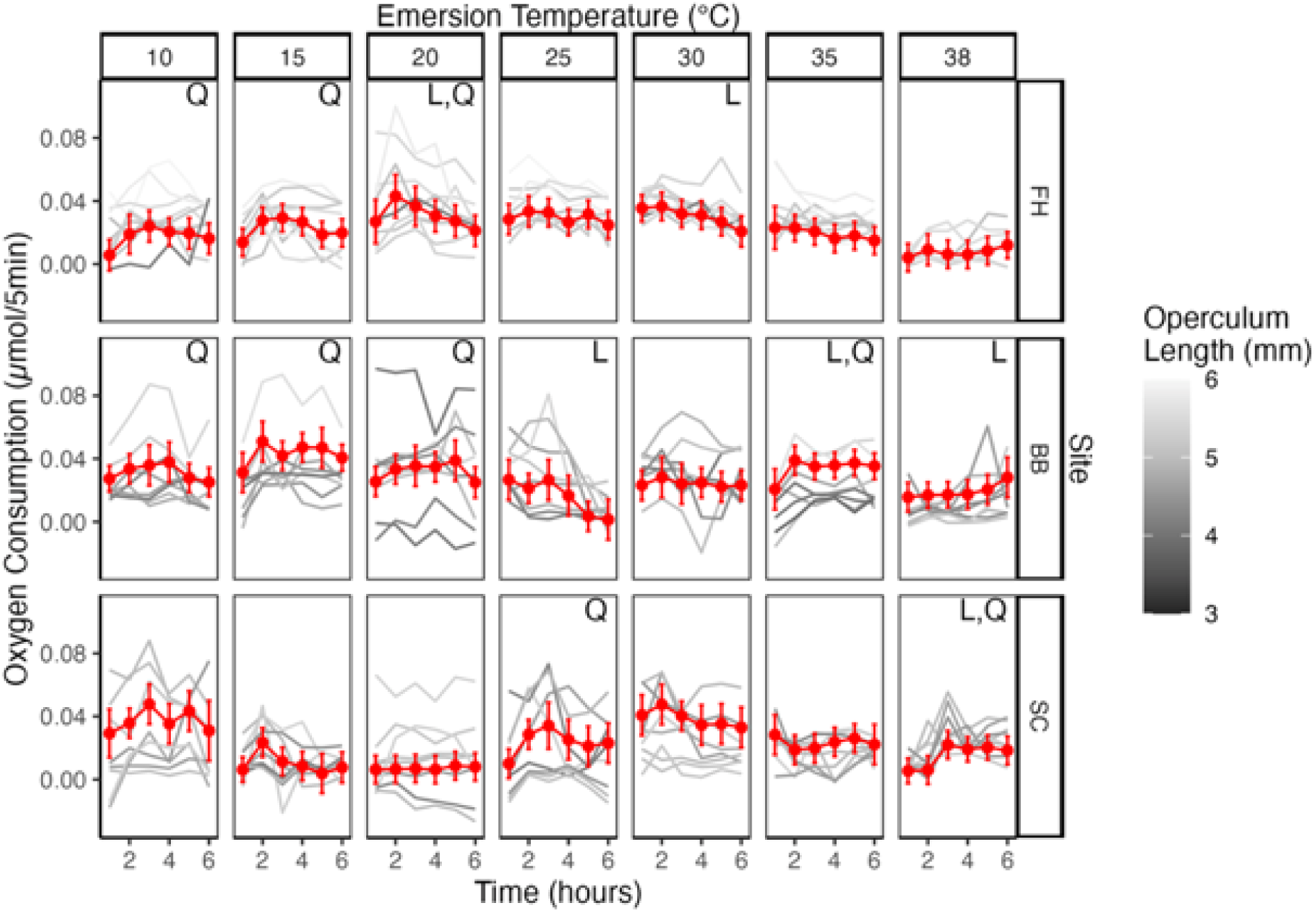
Aquatic oxygen consumption by temperature and hour for each population. Estimated marginal means (±95CI), for a barnacle of 5.05 mm operculum length, are shown in red. Gray lines are data for individual barnacles, with the darkness of the line indicating size. Letters indicate significant polynomial contrasts (p<0.05, L=Linear, Q=Quadratic).

### 3.4 Total cost of emersion and vulnerability to warming

The estimated total cost of a 30-minute emersion ranged from -0.07µmol for BB barnacles at 38°C to 0.66µmol for FH barnacles at 25°C (Fig. 7A). On average, immersion respiration accounted for 18% of the total cost of exposure, but this value ranged from 0 to nearly 50%, depending on the temperature and population.

**Figure 7.**
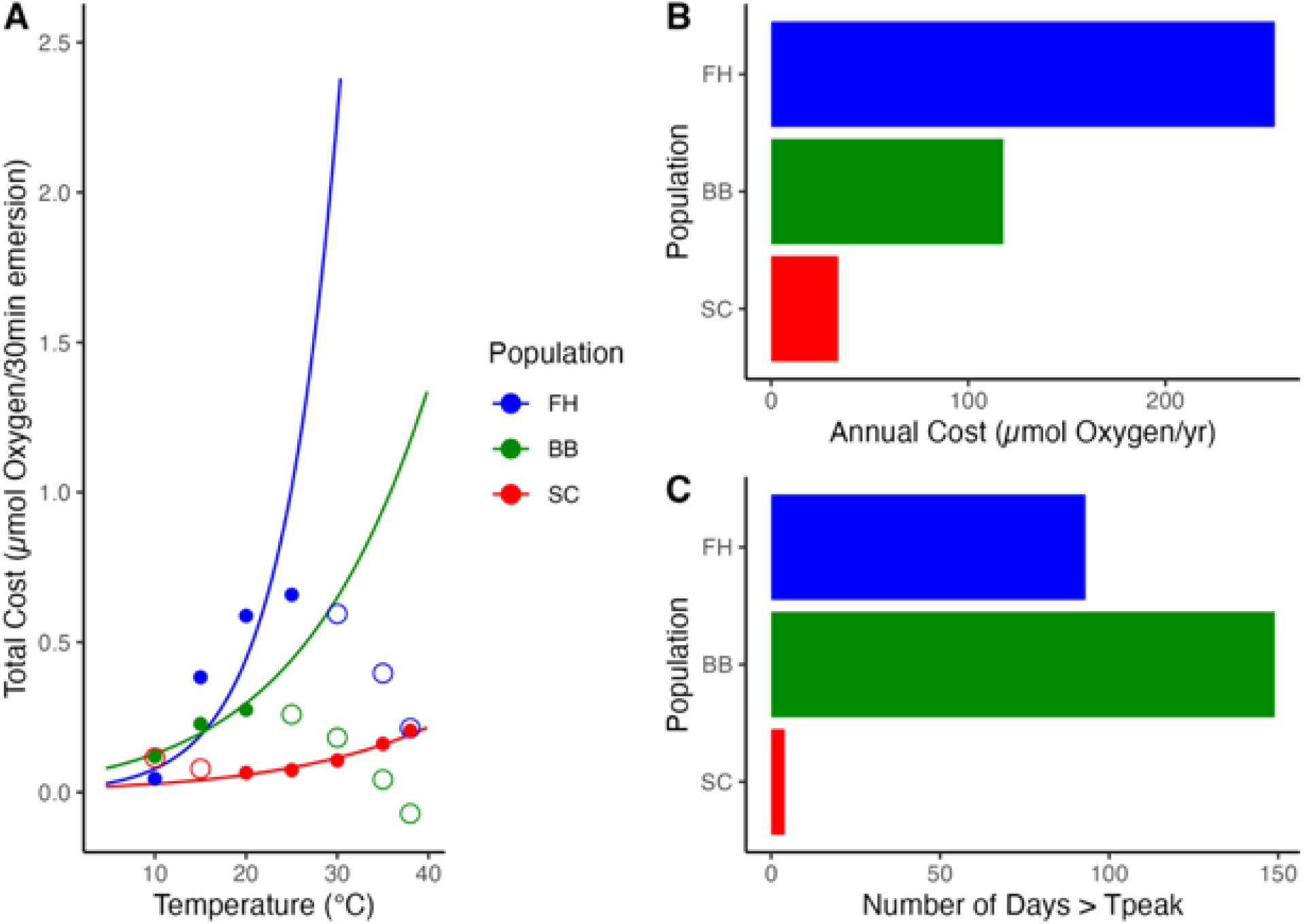
Total cost of a 30-minute emersion exposure by temperature (A), total annual cost for each population (B), and number of days over T_Peak_ (C). Exponential cost curves (solid lines) in (A) are fit on the rising portion of the cost curve (solid dots) and used to calculate total annual costs (B). Both hollow and solid dots in (A) are used to calculate T_peak_ values for (C). See text for additional explanation of cost calculations.

As before, the fitted parabolas for the total cost for the two most northern populations were concave (Table 4), with T_peak_ estimated at 25.1°C (25.12-25.12 95%CI) for FH and 21.3 (21.3-21.3 95%CI) for BB (Fig. 7); while the parabola for SC was convex, with a minimum cost at 20.5°C (20.5-20.5 95%CI, Table 4). The convex shape of the SC curves suggests we may not have tested this population at temperatures above its T_peak_ The MTE-based cost curves showed increasing costs with latitude (Fig. 7A, Table 4). When we these values to calculate the number of days above T_peak_ for each site (Fig. 7C), substituting 38°C for the SC T_peak_, we found BB to be most vulnerable, with 149 days over T_peak_, followed by FH (93 days) and SC (4 days).

**Table 4.** Parameter values (±SE) for the total cost equations fitted for each population.

|  | **Friday Harbor** | **Bodega Bay** | **Southern California** |
| --- | --- | --- | --- |
| **Parabola** |  |  |  |
| Intercept | -1.04 (1.63E-15) | -0.285 (1.63E-15) | 0.257 (1.63E-15) |
| Temperature | 0.135 (1.49E-16) | 0.0529 (1.49E-16) | -0.0187 (1.49E-16) |
| Temperature^2 | -0.00268 (3.04E-18) | -0.00124 (3.04E-18) | 0.000457 (3.04E-18) |
| **Exponential** |  |  |  |
| -E | 48.43 (11.32) | 22.53 (17.74) | 17.77 (8.99) |
| ln($R_0$) | -14434.01 (3287.7) | -6961.4 (5111.17) | -6042.32 (2721.12) |

The exponentially increasing total cost functions (Fig. 7A, solid lines; Table 4) were calculated from the 30-minute emersion costs using only temperatures below (BB, FH) or above (SC) the T_peak_ value. FH had the largest total costs at all temperatures, followed by BB and SC. When we used these equations to calculate the total annual emersion costs for each site (7B), the costs followed the ordering of the cost curves (Fig. 7A), with FH showing the highest energy costs and SC the lowest.

## 4 Discussion

Intertidal organisms experience two very different thermal regimes during emersion and immersion. Understanding the influence of each environment on their thermal physiology is essential for identifying which species and populations are most vulnerable to warming temperatures under climate change. We compared the metabolic costs of emersion for the intertidal barnacle *Balanus glandula* from three populations with contrasting emersion and immersion temperatures. Our results suggest a greater overall influence of local emersion temperatures than immersion temperatures. Specifically, the peaks (T_peak_) of the respiration curves in all three populations exceeded mean aquatic DPM temperatures by 10°C or more and are better matched to the geographic ordering of daily intertidal DPMs. However, this conclusion is tempered by the small thermal differences in intertidal temperatures between our two northernmost sites which limited the statistical significance of some comparisons. Additionally, in using these results to predict the vulnerability of each population to climate change, we found different patterns of vulnerability to warming temperatures depending on the metric of vulnerability used. The most vulnerable site according to the cost approach was the northernmost site, while it was the central site according to the threshold approach. This discrepancy arises from the large differences we observed in energy demand among our three populations, which may reflect local adaptation to nearshore ocean productivity and food supply.

Two lines of evidence in our results suggest that the extreme temperatures experienced during low tide conditions drive the thermal physiology of *B. glandula* during emersion more than immersion temperatures. First, the identified T_peak_ of the total energy cost parabola for each population is much greater than its site’s average aquatic DPM, exceeding these values by at least 10°C at all three sites. While, a population’s T_peak_ should exceed its average DPM to minimize the amount of time spent at stressful temperatures (Martin & Huey 2008), the differences here are much larger than would be expected. Second, the ordering of the thermal peaks of the northern FH and central BB sites match the ordering of emersion DPMs rather than immersion DPMs. Specifically, FH had both warmer emersion temperatures and colder immersion temperatures than BB, but its T_peak_ was warmer than BB. While the temperature differences between the FH and BB sites were small, and not always statistically significant, temperature differences on this scale can greatly influence intertidal community composition and population dynamics (e.g., Barry et al. 1995, Harley 2011).

We also note that the T_peak_ value for the southernmost (SC) population is greater than the highest temperature we tested (38°C), placing it well above the T_peak_ values for BB and FH. We did not expect this result, given that a 40°C emersion is lethal for FH barnacles (Ober et al. 2019). However, it is consistent with the warmer overall temperatures at this site and with previously identified genetic differences among populations of *B. glandula* (Wares & Skoczen 2019, Wares et al. 2021). There is a known thermal and biogeographic breakpoint at Point Conception, California (Wares et al. 2001, Blanchette et al. 2007), which falls between our SC and BB study sites (Fig. 1). Many intertidal species exhibit physiologically distinct populations north and south of this break (e.g., Sagarin & Somero 2006, Gleason & Burton 2013). Unfortunately, because both the aquatic and intertidal thermal environments at the southern SC site are warmer than the two other sites, comparisons with SC do not inform the question of whether aquatic or terrestrial environments are the main driver of these physiological differences.

Our finding that emersion conditions drive thermal physiology at BB and FH matches past studies of thermal physiology among NE Pacific intertidal congeners that occupy different intertidal shore heights. Despite experiencing the same immersion conditions, higher shore congeners typically show a more warm-adapted thermal physiology than their low shore congeners (Somero 2002, 2010). But inter-latitudinal studies of variation in thermal physiology with NE Pacific intertidal species, particularly among sites north of Point Conception, have not consistently shown a role for emersion conditions in thermal physiology (Rao 1953, Sagarin & Somero 2006, Kuo & Sanford 2009, Logan et al. 2012). One explanation for this discrepancy could stem from differences among species in the degree to which they are metabolically active during low tide. *B. glandula* displayed high rates of emersion oxygen consumption, often higher than they exhibited in water at the same temperature (Ober et al. 2019), but many other species exhibit metabolic depression at low tide (e.g., Sokolova & Portner 2001, Marshall et al. 2011). If species are not metabolically active during emersion, then their physiology may be under stronger natural selection during immersion than emersion. Alternately, thermal physiology may vary by trait (Dell et al. 2011, Rose et al. 2024) and if the study measures a trait that is not used during emersion (e.g. feeding rates, locomotion, or pumping rates) then it may only be under natural selection during immersion. In the case of *B. glandula*, however, a separate study of aquatic respiration rates in the FH and LA populations (Ober et al. *unpubl. data*) still found T_peak_ values at water temperatures well over the local maxima.

We used long-duration respiration trials to fully capture the temporal pattern of respiration during emersion and immersion (McGaw et al. 2015, Ober et al. 2019, Griffen et al. 2024). We found significant effects of emersion temperature on the temporal patterns of respiration during both the emersion and immersion phases of the trials. We detected three distinct temporal patterns of respiration during immersion. At the coolest temperatures, barnacles in all three populations exhibited parabolic patterns with initially low respiration rates that increased in the second or third hour of immersion and then declined. We hypothesize that barnacles initially reduced respiration upon immersion because they had access to sufficient oxygen during emersion and were not experiencing high energy demand. Video observations of the FH population (Ober et al. 2019) indicate that *B. glandula* are frequently closed in the first few hours of immersion at these cooler temperatures. Similar patterns of low post-emersion oxygen demand have been reported in other high shore barnacle species (Davenport & Irwin 2003). The concentration of oxygen in air is more than 20-fold higher than in seawater, so oxygen may be more accessible during emersion at non-stressful temperatures then during immersion (Hughes & Otto 1999). By staying closed barnacles may also reduce to vulnerability predation during immersion (Horn et al. 2021).

At warmer temperatures, barnacles began immersion with higher initial respiration rates that declined gradually over the 6-hr immersion period. The high initial immersion respiration coincides with high initial activity in the video data recorded from FH barnacles (2019) and suggests that high energy demand during emersion created an immediate need for respiration upon resubmersion. At these same temperatures, barnacles often also showed a decline in emersion respiration over time, suggesting an oxygen debt was forming (Ellington 1983, McGaw et al. 2015). Finally, at the warmest temperatures, immersion respiration was depressed and remained low for several hours. This third pattern suggests physiological impairment. At these temperatures the barnacles often appeared in a heat coma during immersion with cirri extended but not moving.

The second goal of this study was to assess each population’s vulnerability to warming temperatures under climate change. Surprisingly, our assessment of vulnerability depended on the metric of vulnerability. When we calculated the total annual costs of exposure to intertidal DPM temperatures, the northernmost FH population had the greatest total energy cost among the three sites. However, when we calculated the number of days above T_peak_, the central BB site appeared most vulnerable. We were unable to identify any published studies that compared these two metrics of vulnerability, so it is unclear how common our result is. But it suggests that estimates of vulnerability to warming vary with the metric used.

Greater energetic costs only reflect a vulnerability to warming temperatures if energy intake cannot meet demand. The FH site showed greater overall energy demand at most temperatures, particularly during emersion, which could reflect difference in overall metabolism, driven by greater food availability at higher latitudes. *Balanus glandula*’s exact diet is unknown (Sanford & Menge 2001), but they likely consume a range of phytoplankton and zooplankton, as adults can thrive on either in the laboratory (Hines 1978, Geierman & Emlet 2009, Gilman et al. 2013). There is a strong latitudinal gradient in primary productivity throughout this region of the northeastern Pacific, with coastal Washington State and British Columbia exhibiting 5-fold greater chlorophyll concentrations than California (Ware & Thomson 2005, Hickey & Banas 2008), along with higher zooplankton concentrations (Ware & Thomson 2005). This gradient is attributed to greater terrestrial nutrient supply at higher latitudes, particularly in the Salish Sea region (Hickey & Banas 2008), which includes the FH field site. A greater food supply could select for greater overall energy use in the two higher latitude populations, while the SC (southernmost) barnacles may be adapted to succeed in a lower resource environment. This is an alternate explanation for previously described genetic cline in *B. glandula* (Sotka et al. 2004, Wares et al. 2021). If true, this suggests that energy costs are a less useful metric of vulnerability to warming than threshold temperatures for NE Pacific suspension feeders such as *B. glandula*.

## 5 Conclusion

The results presented herein provide strong evidence that the thermal conditions experienced during emersion influence the thermal physiology of the intertidal barnacle *Balanus glandula*. However, more empirical studies are needed to predict vulnerability to warming temperatures under climate change. The possibility that *B. glandula* populations are locally adapted or acclimated to both thermal regimes and food supplies complicates predictions of their response to warming temperatures. While its well-established that food supply can limit many ectotherms’ tolerance of thermal stress (e.g., Gilman 2006, Schneider et al. 2010), few models of climate change responses have considered both warming temperatures and changes in food supply at the same time (but see Huey & Kingsolver 2019). While its higher overall energy demand suggests that the FH population is the most vulnerable to warming temperatures, phytoplankton productivity is also predicted to increase in this region under climate change (Khangaonkar et al. 2019). Thus, in this case, the central BB population may be the most vulnerable to future warming under climate change.

## Acknowledgements

This work was supported by the National Science Foundation (grant IOS-1351445 to S.E.G.) with additional support from the Keck Science Department of Claremont McKenna, Pitzer, and Scripps Colleges. We thank E. Roberts for commenting on an earlier draft of this manuscript. The present work is available as preprint at https://www.biorxiv.org/content/10.1101/2024.03.26.586848v1.

